## Supplemental figures and table for "Modelling group heteroscedasticity in single-cell RNA-seq pseudo-bulk data"

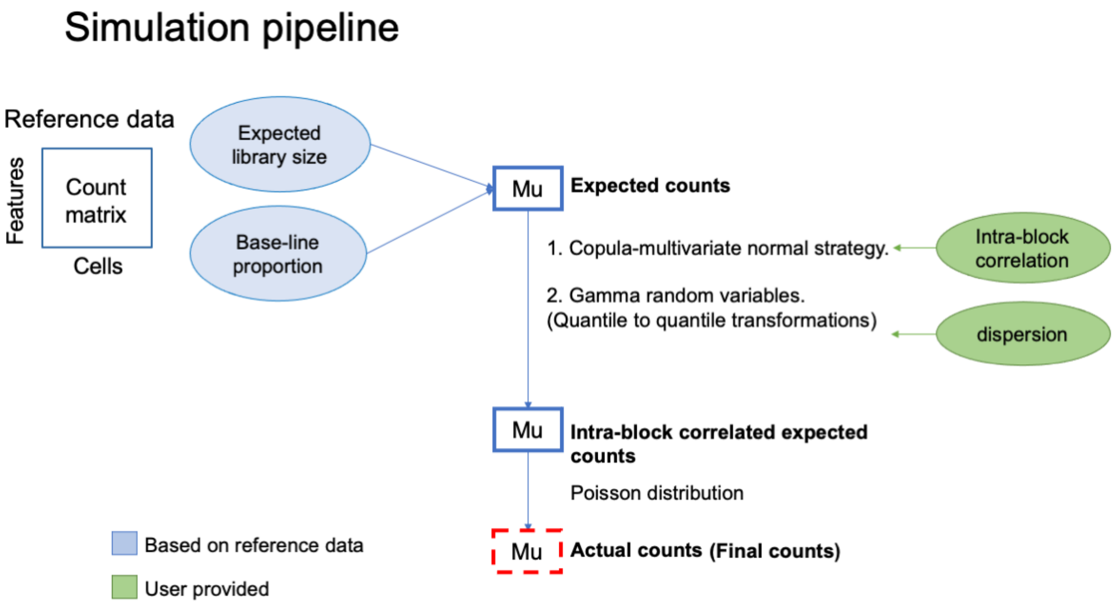


Supplementary Figure S1, Simulation pipeline to generate scRNA-seq data. Expected library size and baseline proportions are estimated on reference scRNA-seq data to get an expected count matrix. Copula-multivariate normal strategy is used to simulate intra-block correlation between cells. Then they are transformed to be Gamma random variables. The final counts are generated from a Poisson distribution.

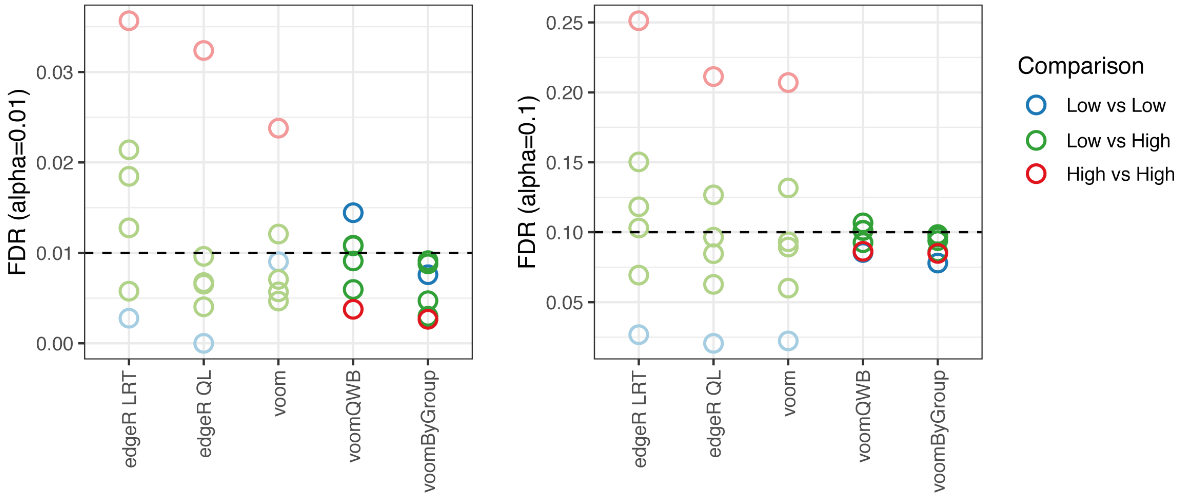


Supplementary Figure S2, Comparing FDR across DE methods at cut-off set as 0.01 and 0.1 under the scenario with unequal group biological variation. Color denotes comparison type.

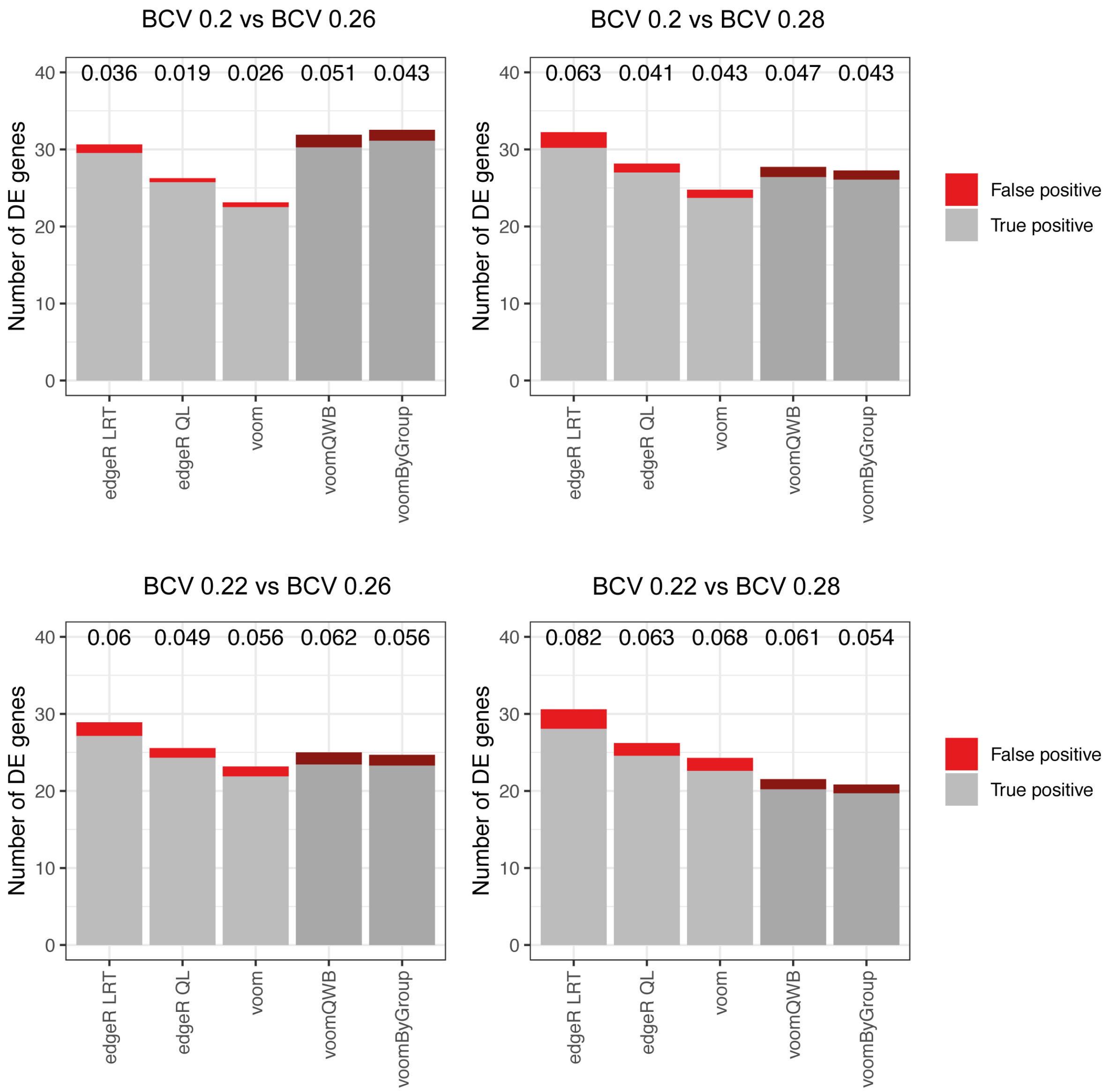


Supplementary Figure S3, Comparing the power of DE methods to detect true DE genes under the scenario with unequal group biological variation. In each comparison, true positive genes are plotted in grey, and false-positive genes are plotted in red. FDR by each of the methods is labelled at the top of the bar.

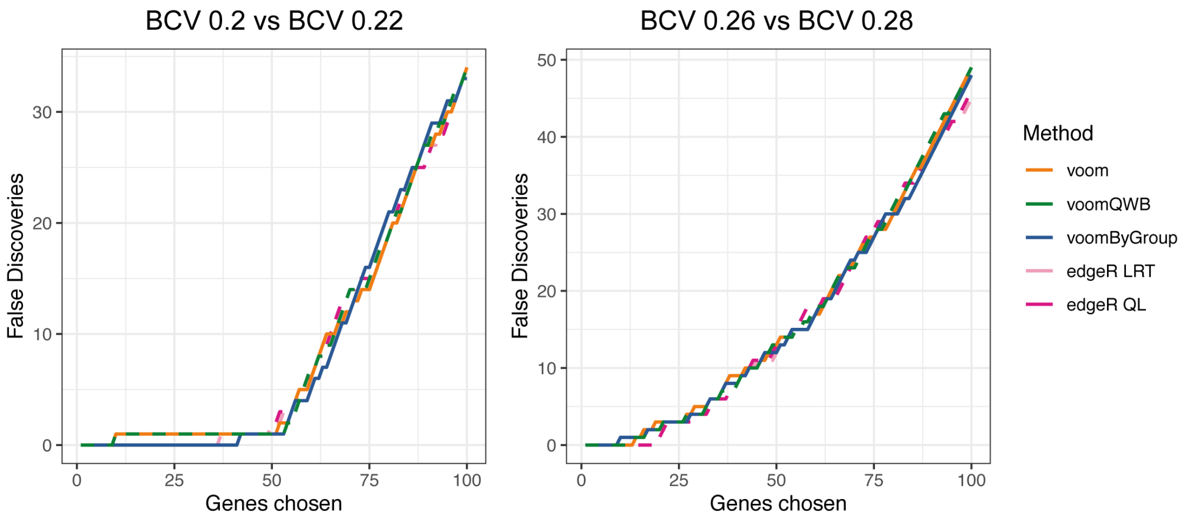


Supplementary Figure S4, Cumulative false discoveries across 50 simulated data sets under the scenario with unequal group biological variation. The y axis shows the cumulative number of false positives amongst the top 100 genes from each comparison. Color denotes the DE method.

a BCV 0.26 vs BCV 0.28

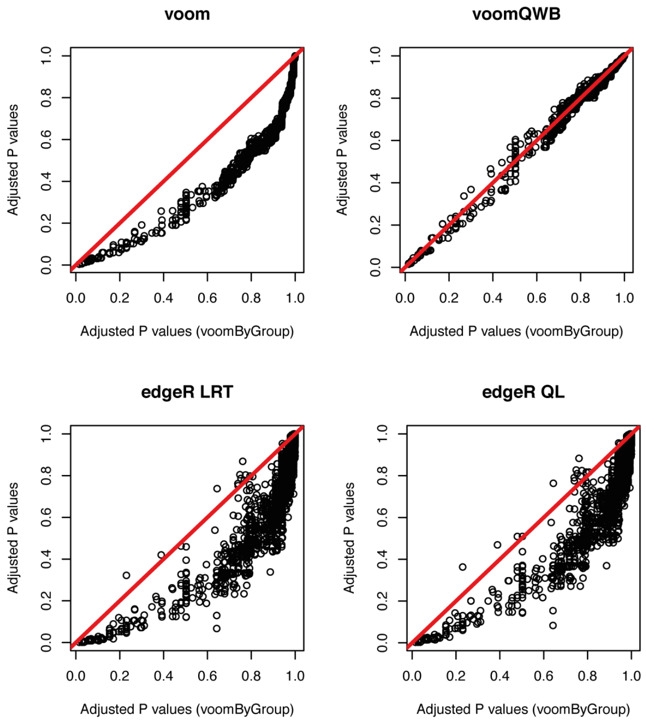


b BCV 0.2 vs BCV 0.22

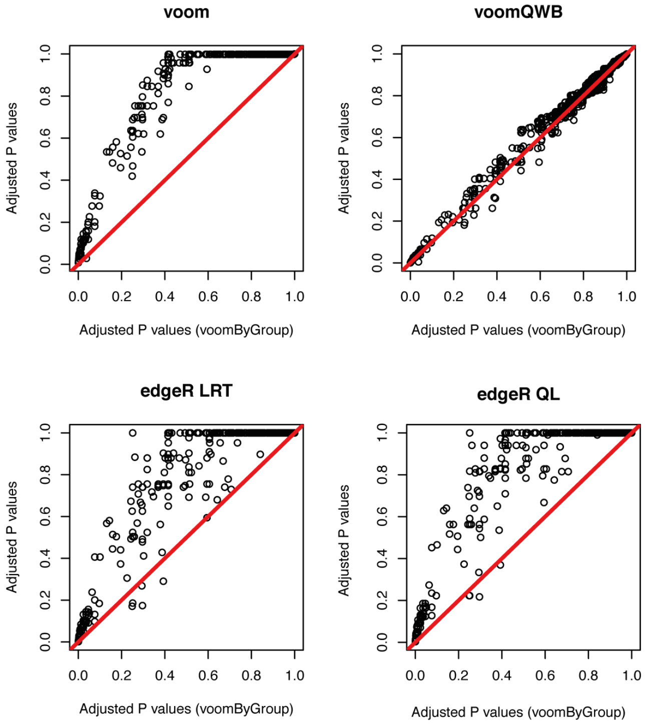


Supplementary Figure S5, Pair-wise comparisons between adjusted P values from *voomByGroup* to adjusted P values from other methods in a) comparison of high-BCV group compared to high-BCV group, and b) comparison of low-BCV group compared to low-BCV group. y=x is plotted in the red line.

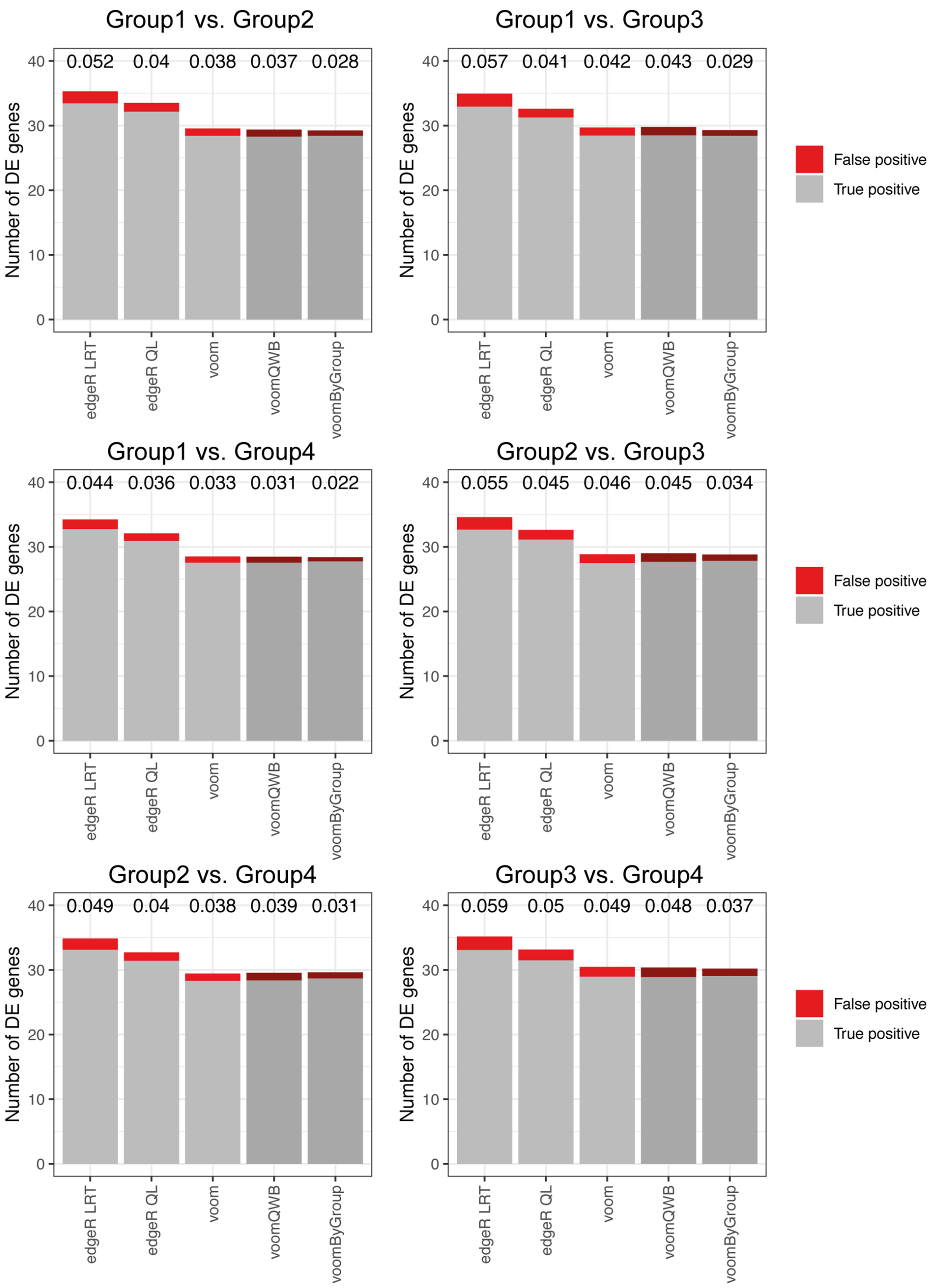


Supplementary Figure S6, Comparing the power of DE methods to detect true DE genes in the null simulation. In each comparison, true positive genes are plotted in grey, and false-positive genes are plotted in red. FDR by each of the methods is labelled at the top of the bar.

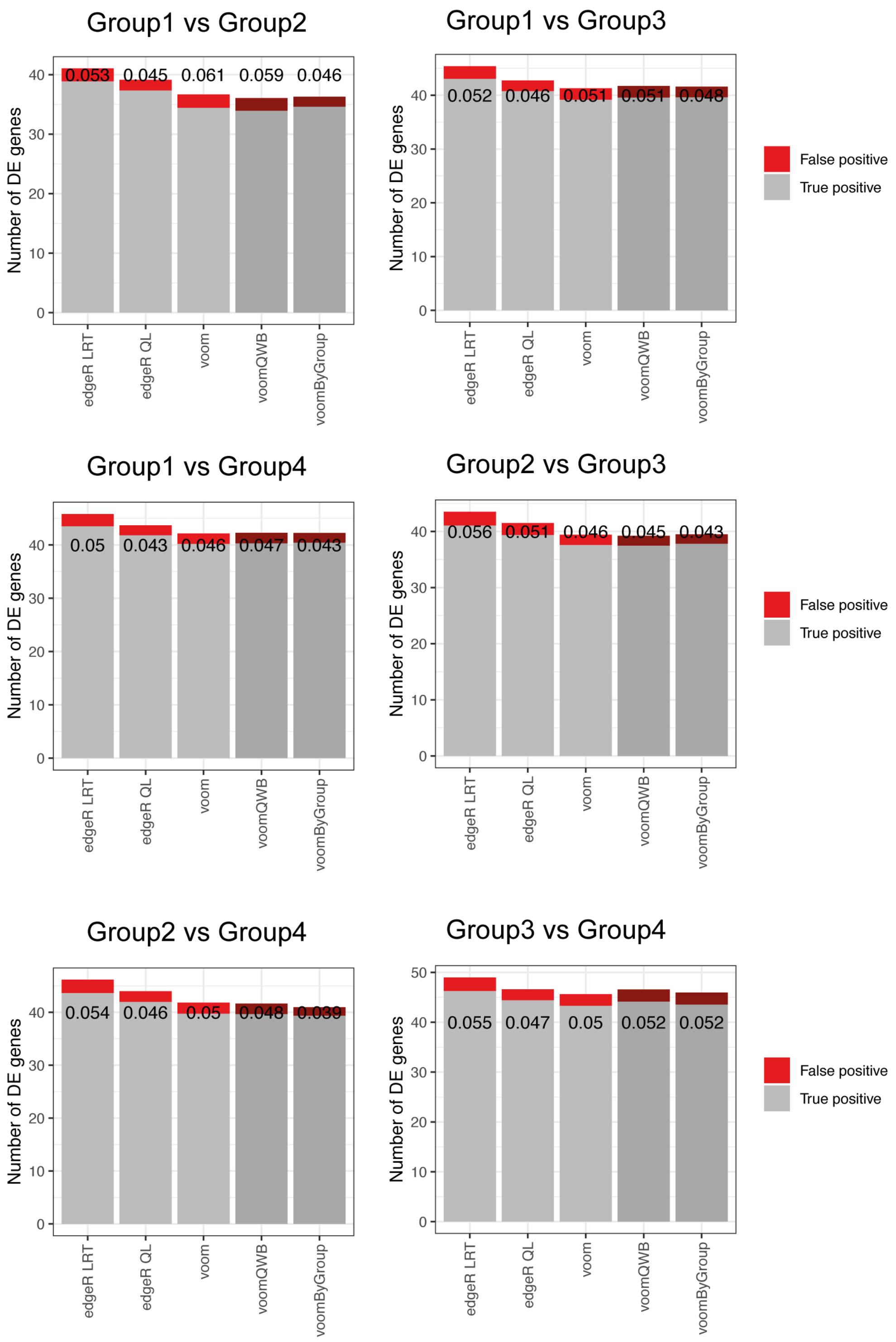


Supplementary Figure S7, Under the scenario with unequal sample sizes, at cut-off 0.05, the number of DE genes delivered by different methods is plotted across different comparisons. In the bar plot, grey represents true positive genes, and red represents false positive genes. FDR by each of the method is labelled at the top of the bar.

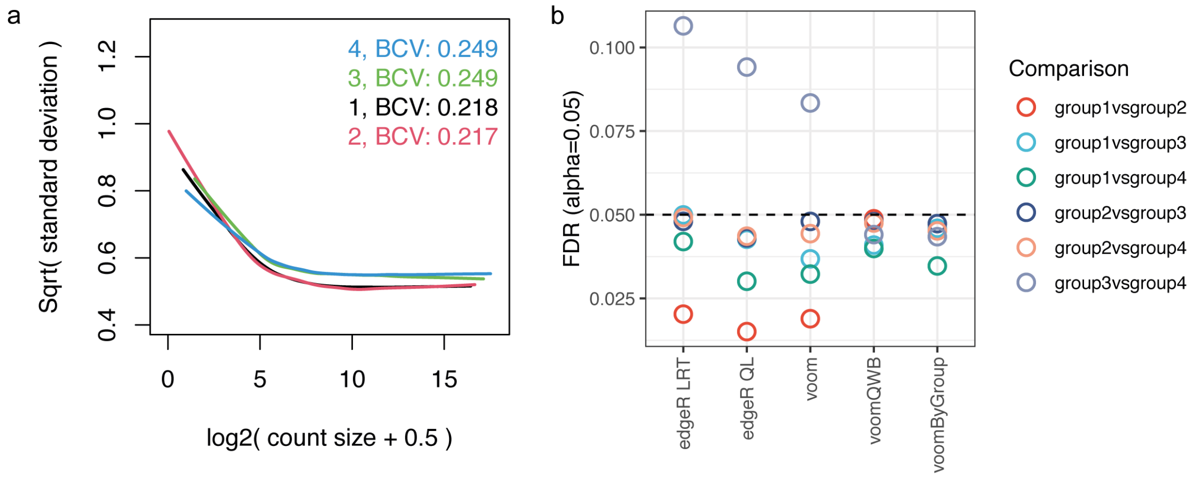


Supplementary Figure S8, Under the scenario with unequal sample sizes and unequal group-wise BCV, group-wise mean-variance trends are plotted in a). FDRs across methods in different comparisons (colors denote comparisons) are summarized in b) given the cut-off set as 0.05.

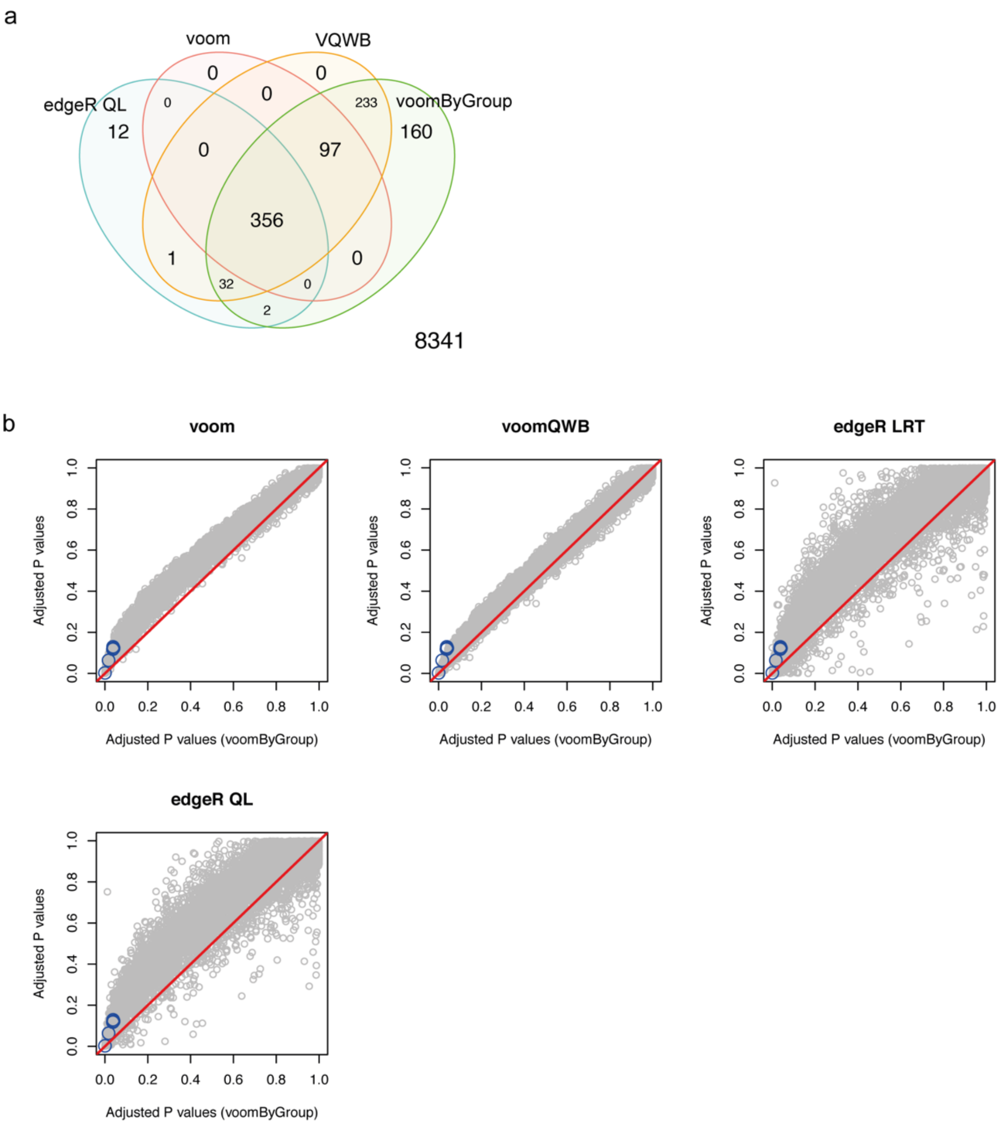


Supplementary Figure S9, Performance of DE methods comparing CD56$^{dim}$CD16$^{+}$ NK cells between asymptomatic patients to healthy control.
Venn diagram showing the number of genes DE in the comparison between asymptomatic patients and healthy controls delivered by edgeR (quasi-likelihood), *voom*, *voomQW*, *voomByGroup* from left to right. The number of genes that are not DE in either comparison is labelled in the bottom-right. Pair-wise P values comparisons are shown in b) with scatter plot. Y=x is plotted in the red line. Genes detected to be interferon-gamma related by *voomByGroup* are colored in blue.

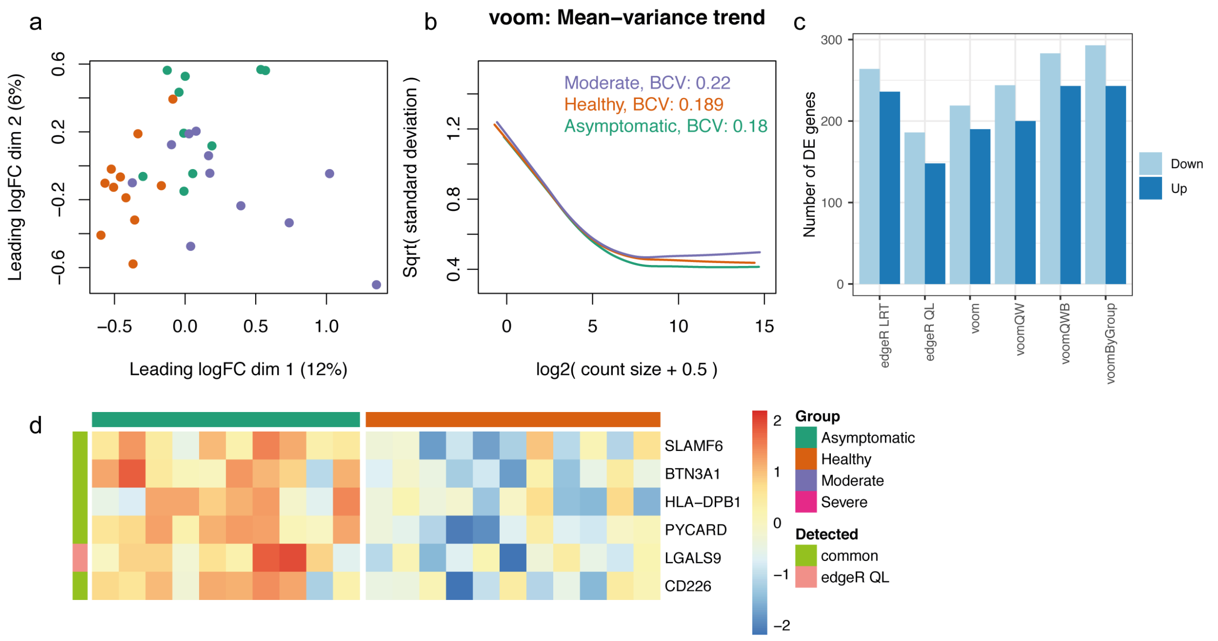


Supplementary Figure S10, DE analysis on dataset PBMC2. Multidimensional scaling plot of $CD56^{dim}CD16^{+}$ NK cells' pseudo-bulk samples is plotted in a), with colors denoting conditions. Mean-variance trends in the individual condition are plotted in b). The number of DE genes is shown in the bar plot in c) with colors denoting up- or down- regulation. Expression of commonly detected up-regulated DE genes and one uniquely detected gene by *edgeR QL* in asymptomatic patients are shown in d).

**Supplementary Table S1: Summary of filtering steps applied to each dataset**

| Dataset | Gene filtering | Cell filtering | Selected cell type | Pseudo-bulk sample filtering |
|---|---|---|---|---|
| Simulated data | total counts < 30 removed | - | - | - |
| Mouse lung data | total counts < 200 removed | total counts < 200 removed | type2 pneumocytes | Number of cells < 50 removed |
| Xenopus tail data | total counts < 50 removed | total counts < 200 removed | goblet cells | Number of cells < 40 removed |
| Human lung data | total counts < 200 removed | total counts < 200 removed | macrophages | Number of cells < 80 removed |
| Human PBMC COVID data 1 | total counts < 50 removed | total counts < 200 removed | CD56dim CD16+ NK cells | Number of cells < 50 removed |
| Human PBMC COVID data 2 | total counts < 50 removed | total counts < 200 removed | CD56dim CD16+ NK cells | Number of cells < 50 removed & Library sizes < 200,000 removed |
